# Immune-related genetic enrichment in frontotemporal dementia

**DOI:** 10.1101/157875

**Authors:** Iris Broce, Celeste M. Karch, Natalie Wen, Chun C. Fan, Yunpeng Wang, Chin Hong Tan, Naomi Kouri, Owen A. Ross, Günter U. Höglinger, Ulrich Muller, John Hardy, International FTD-Genomics Consortium (IFGC), Parastoo Momeni, Christopher P. Hess, William P. Dillon, Zachary A. Miller, Luke W. Bonham, Gil D. Rabinovici, Howard J. Rosen, Gerard D. Schellenberg, Andre Franke, Tom H. Karlsen, Jan H. Veldink, Raffaele Ferrari, Jennifer S. Yokoyama, Bruce L. Miller, Ole A. Andreassen, Anders M. Dale, Rahul S. Desikan, Leo P. Sugrue, R Ferrari, D G Hernandez, M A Nalls, J D Rohrer, A Ramasamy, J B J Kwok, C Dobson-Stone, P R Schofield, G M Halliday, J R Hodges, O Piguet, L Bartley, E Thompson, E Haan, I Hernández, A Ruiz, M Boada, B Borroni, A Padovani, C Cruchaga, N J Cairns, L Benussi, G Binetti, R Ghidoni, G Forloni, D Albani, D Galimberti, C Fenoglio, M Serpente, E Scarpini, J Clarimón, A Lleó, R Blesa, M Landqvist Waldö, K Nilsson, C Nilsson, I R A Mackenzie, G-Y R Hsiung, D M A Mann, J Grafman, C M Morris, J Attems, T D Griffiths, I G McKeith, A J Thomas, P Pietrini, E D Huey, E M Wassermann, A Baborie, E Jaros, M C Tierney, P Pastor, C Razquin, S Ortega-Cubero, E Alonso, R Perneczky, J Diehl-Schmid, P Alexopoulos, A Kurz, I Rainero, E Rubino, L Pinessi, E Rogaeva, P St George-Hyslop, G Rossi, F Tagliavini, G Giaccone, J B Rowe, J C M Schlachetzki, J Uphill, J Collinge, S Mead, A Danek, V M Van Deerlin, M Grossman, J Q Trojanowski, J van der Zee, M Cruts, C Van Broeckhoven, S F Cappa, I Leber, D Hannequin, V Golfier, M Vercelletto, A Brice, B Nacmias, S Sorbi, S Bagnoli, I Piaceri, J E Nielsen, L E Hjermind, M Riemenschneider, M Mayhaus, B Ibach, G Gasparoni, S Pichler, W Gu, M N Rossor, N C Fox, J D Warren, M G Spillantini, H R Morris, P Rizzu, P Heutink, J S Snowden, S Rollinson, A Richardson, A Gerhard, A C Bruni, R Maletta, F Frangipane, C Cupidi, L Bernardi, M Anfossi, M Gallo, M E Conidi, N Smirne, R Rademakers, M Baker, D W Dickson, N R Graff-Radford, R C Petersen, D Knopman, K A Josephs, B F Boeve, J E Parisi, W W Seeley, B L Miller, A M Karydas, H Rosen, J C van Swieten, E G P Dopper, H Seelaar, Y A L Pijnenburg, P Scheltens, G Logroscino, R Capozzo, V Novelli, A A Puca, M Franceschi, A Postiglione, G Milan, P Sorrentino, M Kristiansen, H-H Chiang, C Graff, F Pasquier, A Rollin, V Deramecourt, T Lebouvier, D Kapogiannis, L Ferrucci, S Pickering-Brown, A B Singleton, J Hardy, P Momeni

## Abstract

**Background:** Converging evidence suggests that immune-mediated dysfunction plays an important role in the pathogenesis of frontotemporal dementia (FTD). Although genetic studies have shown that immune-associated loci are associated with increased FTD risk, a systematic investigation of genetic overlap between immune-mediated diseases and the spectrum of FTD-related disorders has not been performed.

**Methods and findings:** Using large genome-wide association studies (GWAS) (total n = 192,886 cases and controls) and recently developed tools to quantify genetic overlap/pleiotropy, we systematically identified single nucleotide polymorphisms (SNPs) *jointly* associated with ‘FTD-related disorders’ namely FTD, corticobasal degeneration (CBD), progressive supranuclear palsy (PSP), and amyotrophic lateral sclerosis (ALS) – and one or more immune-mediated diseases including Crohn’s disease (CD), ulcerative colitis (UC), rheumatoid arthritis (RA), type 1 diabetes (T1D), celiac disease (CeD), and psoriasis (PSOR). We found up to 270-fold genetic enrichment between FTD and RA and comparable enrichment between FTD and UC, T1D, and CeD. In contrast, we found only modest genetic enrichment between any of the immune-mediated diseases and CBD, PSP or ALS. At a conjunction false discovery rate (FDR) < 0.05, we identified numerous FTD-immune pleiotropic SNPs within the human leukocyte antigen (*HLA)* region on chromosome 6. By leveraging the immune diseases, we also found novel FTD susceptibility loci within *LRRK2* (Leucine Rich Repeat Kinase 2)*, TBKBP1* (TANK-binding kinase 1 Binding Protein 1), and *PGBD5* (PiggyBac Transposable Element Derived 5). Functionally, we found that expression of FTD-immune pleiotropic genes (particularly within the *HLA* region) is altered in postmortem brain tissue from patients with frontotemporal dementia and is enriched in microglia compared to other central nervous system (CNS) cell types.

**Conclusions:** We show considerable immune-mediated genetic enrichment specifically in FTD, particularly within the *HLA* region. Our genetic results suggest that for a subset of patients, immune dysfunction may contribute to risk for FTD. These findings have potential implications for clinical trials targeting immune dysfunction in patients with FTD.

## INTRODUCTION

Frontotemporal dementia (FTD) is a fatal neurodegenerative disorder and the leading cause of dementia among individuals younger than 65 years of age [1]. Given rapid disease progression and absence of disease modifying therapies, there is an urgent need to better understand FTD pathobiology to accelerate development of novel preventive and therapeutic strategies.

Converging molecular, cellular, genetic, and clinical evidence suggests that neuroinflammation plays a role in FTD pathogenesis. Complement factors and activated microglia, key components of inflammation, have been established as histopathologic features in brains of patients [2] and in mouse models of FTD [3,4]. Genome-wide association studies (GWAS) have shown that single nucleotide polymorphisms (SNPs) within the immune-regulating human leukocyte antigen (*HLA*) region on chromosome 6 are associated with elevated FTD risk [5]. Importantly, there is increased prevalence of immune-mediated diseases among patients with FTD [6,7]. Together, these findings suggest that immune-related mechanisms may contribute to and potentially drive FTD pathology.

Recent work in human molecular genetics has emphasized ‘pleiotropy’, where variations in a single gene can affect multiple, seemingly unrelated phenotypes [8]. In the present study, we systematically evaluated genetic pleiotropy between FTD and immune-mediated diseases. Leveraging large neurodegenerative GWASs and recently developed tools to estimate polygenic pleiotropy, we sought to identify SNPs *jointly* associated with ‘FTD-related disorders’ [9,10] – namely FTD, corticobasal degeneration (CBD),progressive supranuclear palsy (PSP), and amyotrophic lateral sclerosis (ALS) – and one or more immune-mediated diseases including Crohn’s disease (CD), ulcerative colitis (UC), rheumatoid arthritis (RA), type 1 diabetes (T1D), celiac disease (CeD), and psoriasis (PSOR).

## METHODS

### Participant samples

We evaluated complete GWAS results in the form of summary statistics (p-values and odds ratios) for FTD, CBD, PSP, and ALS and 6 immune-mediated diseases, including CD [11], UC [12], RA [13], T1D [14], CeD [15], and PSOR [16] (see Table 1).We obtained FTD GWAS summary statistic data from phase I of the International FTD-Genomics Consortium (IFGC), which consisted of 2,154 clinical FTD cases and 4,308 controls with genotyped and imputed data at 6,026,384 SNPs (Table 1, for additional details, see [5]). The FTD dataset included multiple clinically diagnosed FTD subtypes: behavioral variant (bvFTD), semantic dementia (sdFTD), primary nonfluent progressive aphasia (pnfaFTD), and FTD overlapping with motor neuron disease (mndFTD). These FTD cases and controls were recruited from forty-four international research groups and diagnosed according to the Neary criteria [17]. We obtained CBD GWAS summary statistic data from 152 CBD cases and 3,311 controls at 533,898 SNPs (Table 1, for additional details see [18]). The CBD cases were collected from eight institutions and controls were recruited from the Children's Hospital of Philadelphia Health Care Network. CBD was neuropathologically diagnosed using the National Institute of Health Office of Rare Diseases Research criteria [19]. We obtained PSP GWAS summary statistic data (stage 1) from the NIA Genetics of Alzheimer’s Disease Storage Site (NIAGADS), which consisted of 1,114 individuals with autopsy-confirmed PSP and 3,247 controls at 531,451 SNPs (Table 1, for additional details see [20]). We obtained publicly-available ALS GWAS summary statistic data from 12,577 ALS cases and 23,475 controls at 18,741,501 SNPs (Table 1, for additional details see [21]). The relevant institutional review boards or ethics committees approved the research protocol of the individual GWAS used in the current analysis, and all human participants gave written informed consent.

**Table 1.** Summary data from all GWAS used in the current study

| **Disease/Trait** |  | **Total N** | **#** | **Reference** |
| --- | --- | --- | --- | --- |
| Frontotemporal Dementia | FTD | 6,462 | 6,026,384 | [5] |
| Corticobasal degeneration | CBD | 3,463 | 533,898 | [18] |
| Progressive Supranuclear Palsy | PSP | 4,361 | 531,451 | [20] |
| Amyotrophic Lateral Sclerosis | ALS | 36,052 | 18,741,501 | [21] |
| Crohn Disease | CD | 51,109 | 942,858 | [11] |
| Ulcerative colitis | UC | 26,405 | 1,273,589 | [12] |
| Rheumatoid arthritis | RA | 25,708 | 2,554,714 | [13] |
| Type 1 diabetes | T1D | 16,559 | 841,622 | [14] |
| Celiac disease | CeD | 15,283 | 528,969 | [15] |
| Psoriasis | PSOR | 7,484 | 1,121,166 | [16] |

### Genetic Enrichment Statistical Analyses

To identify specific loci jointly involved with each of the four FTD-related disorders and the six immune-mediated diseases, we computed conjunction FDRs [22,23]. Conjunction FDR, denoted by FDR _trait1_&_trait2_ is defined as the posterior probability that a SNP is null for either trait or for both simultaneously, given the *p*-values for both traits are as small, or smaller, than the observed *p*-values. Unlike the *conditional* FDR which ranks disease/primary phenotype associated SNPs based on genetic ‘relatedness’ with secondary phenotypes [24], the *conjunction* FDR minimizes the possibility/likelihood of a single phenotype driving the common association signal. *Conjunction* FDR therefore tends to be more conservative and specifically pinpoints pleiotropic loci between the traits/diseases of interest. We used an overall FDR threshold of < 0.05, which means 5 expected false discoveries per hundred reported. We constructed Manhattan plots based on the ranking of conjunction FDR to illustrate the genomic location of the pleiotropic loci. Detailed information on conjunction Q-Q plots, Manhattan plots, and conjunction FDR can be found in prior reports [22,23,25,26].

### Functional evaluation of shared risk loci

To assess whether SNPs that are shared between FTD and immune-mediated disease modify gene expression, we identified *cis*-expression quantitative loci (eQTLs, defined as variants within 1 Mb of a gene's transcription start site) associated with shared FTD-immune SNPs and measured their regional brain expression in a publicly available dataset of normal control brains (UKBEC, http://braineac.org/) [27]. We also evaluated eQTLs using a blood-based dataset [28]. We applied an analysis of covariance (ANCOVA) to test for associations between genotypes and gene expression. We tested SNPs using an additive model.

### Network based functional association analyses

To evaluate potential protein and genetic interactions, co-expression, co-localization, and protein domain similarity for the functionally expressed (i.e. with significant *cis*-eQTLs) overlapping genes, we used GeneMANIA (www.genemania.org),an online web-portal for bioinformatic assessment of gene networks [29]. In addition to visualizing the composite gene network, we also assessed the weights of individual components within the network [30].

### Gene expression alterations in FTD brains

To determine whether functionally expressed (i.e. with significant *cis*-eQTLs) pleiotropic genes are differentially expressed in FTD brains, we analyzed gene expression of overlapping genes in a publically available dataset [31]. Specifically, we analyzed gene expression data from the frontal cortex, hippocampus, and cerebellum of controls and patients with frontotemporal dementia (FTD-U with or without *progranulin* (*GRN*) mutations, total n =28) (Gene Expression Omnibus (GEO) dataset GSE13162) [31].

### Evaluation of cell classes within the brain

Using a publicly available RNA-sequencing transcriptome and splicing database [32], we ascertained whether the functionally expressed (i.e. with significant *cis*-eQTLs) pleiotropic genes are expressed by specific cell classes within the brain. The eight cell types surveyed are neurons, astrocytes, oligodendrocyte precursor cells, newly formed oligodendrocytes, myelinating oligodendrocytes, microglia, endothelial cells, and pericytes (for additional details, see [32]).

## RESULTS

### Shared genetic risk between FTD and immune-mediated disease

Using progressively stringent *p*-value thresholds for FTD SNPs (i.e. increasing values of nominal –log_10_(*p*)), we observed considerable genetic enrichment for FTD as a function of several immune-mediated diseases (Figure 1A). More specifically, we found strong (up to 270-fold) genetic enrichment between FTD and RA, and comparable enrichment between FTD and UC, T1D, and CeD, with weaker enrichment for PSOR and CD.

**Figure 1A.**
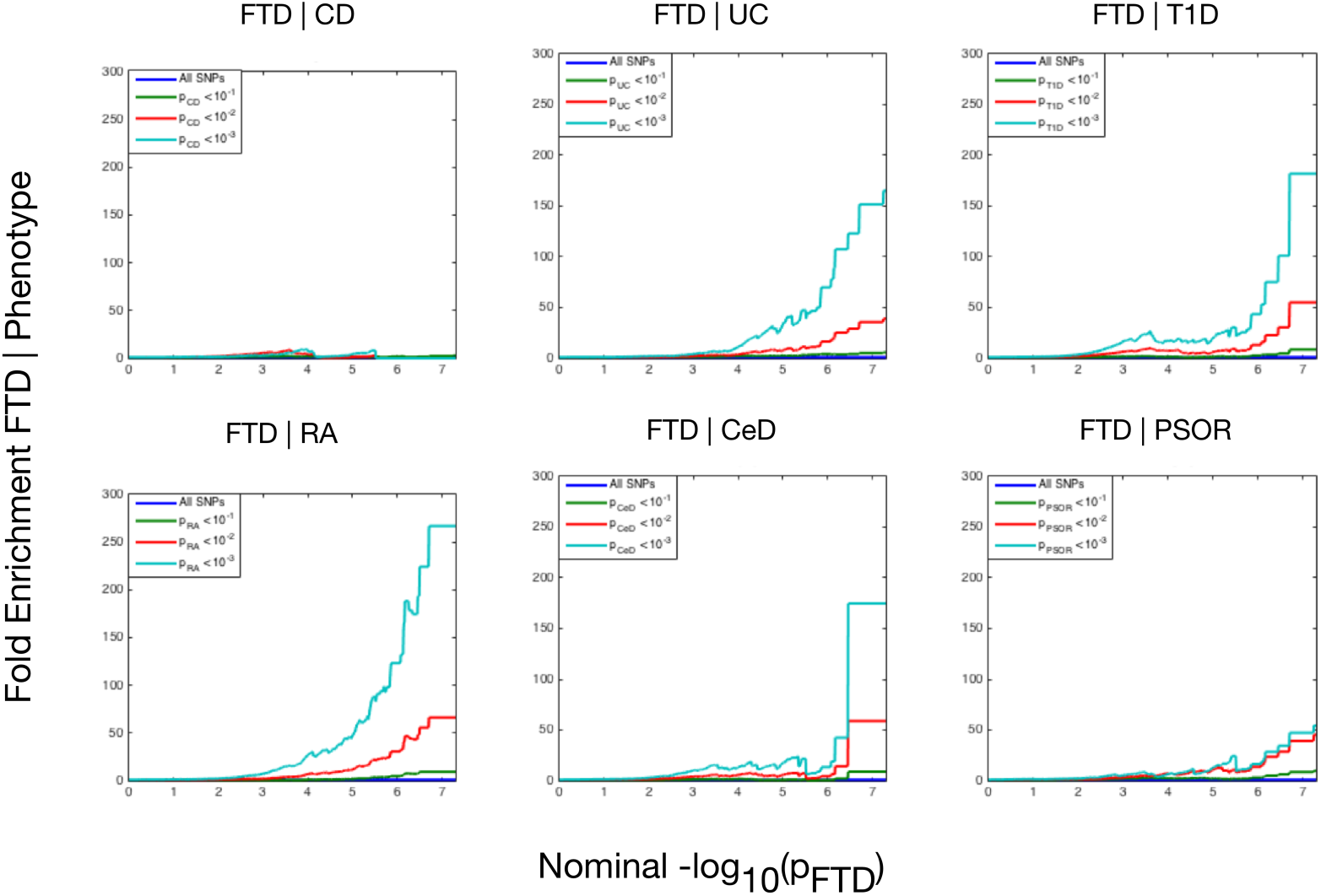
Fold enrichment plots of enrichment versus nominal -log_10_ *p*-values (corrected for inflation) in frontotemporal dementia (FTD) below the standard GWAS threshold of *p* < 5× 10^-8^ as a function of significance of association with 6 immune-mediated diseases, namely Crohn’s disease (CD), ulcerative colitis (UC), type 1 diabetes (T1D), rheumatoid arthritis (RA), celiac disease (CeD), and psoriasis (PSOR) and at the level of -log_10_(*p*) ≥ 0, -log_10_(*p*) ≥ 1, -log_10_(*p*) ≥ 2 corresponding to *p* ≤ 1, *p* ≤ 0.1, *p* ≤ 0.01, respectively. Teal line indicates all SNPs.

At a conjunction FDR *p <* 0.05, we identified 21 SNPs that were associated with both FTD and immune-mediated diseases (Figure 1B, Table 2). Fourteen of these SNPs mapped to the *HLA* region on chromosome 6 (Figure 1B). Of these, three pairs of SNPs showed strong linkage disequilibrium (LD), suggesting that they reflected the same signal: a) rs9261536 with rs3094138 (nearest genes: *TRIM15* and *TRIM26*, respectively, pairwise D’ = 0.96, r^2^ = 0.8), b) rs204991 with rs204989 (nearest gene: *GPSM3*, pairwise D’ = 1, r^2^ = 1), and c) rs9268877 with rs9268852 (nearest gene: *HLA-DRA*, pairwise D’ = 1, r^2^ = 0.99). After excluding those identified SNPs in LD, we found that 11 of the 18 identified loci mapped to the *HLA* region, suggesting that *HLA* markers were critical in driving our results. To test this hypothesis, we repeated our enrichment analysis after removing all SNPs in LD with r^2^ > 0.2 within 1Mb of *HLA* variants (based on 1000 Genomes Project LD structure). After removing *HLA* SNPs, we saw considerable attenuation of genetic enrichment in FTD as a function of immune-mediated disease (Figure 2), suggesting that the observed overlap between immune-related diseases and FTD was largely driven by the *HLA* region.

**Figure 1B.**
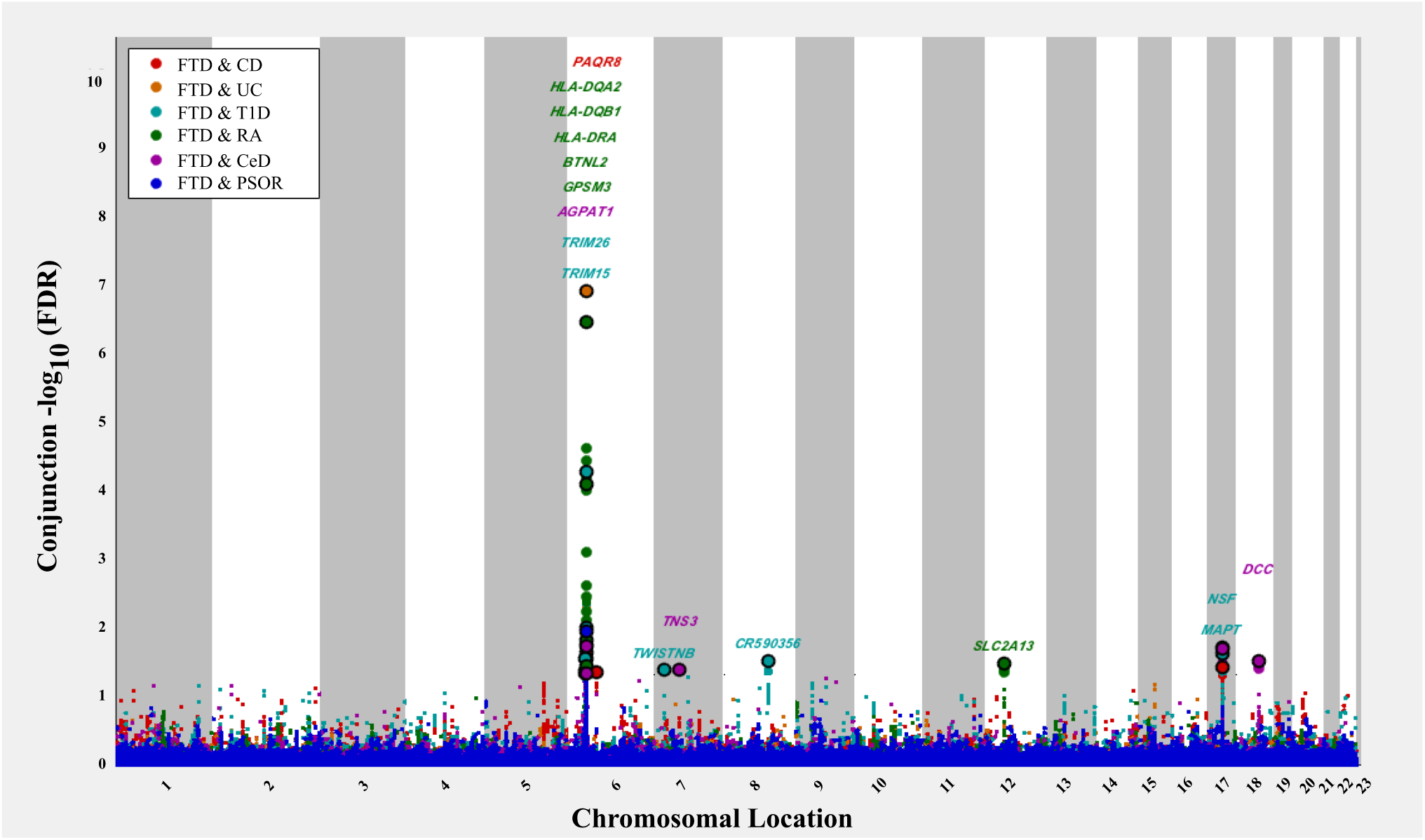
‘Conjunction’ Manhattan plot of conjunction –log_10_ (FDR) values for frontotemporal dementia (FTD) given Crohn’s disease (CD; FTD|CD, red), ulcerative colitis (UC, FTD|UC, orange), type 1 diabetes (T1D, FTD|T1D, teal), rheumatoid arthritis (RA, FTD|RA, green), celiac disease (CeD, FTD|CeD, magenta), and psoriasis (PSOR, FTD|PSOR, blue). SNPs with conjunction –log_10_ FDR > 1.3 (i.e. FDR < 0.05) are shown with large points. A black line around the large points indicates the most significant SNP in each LD block and this SNP was annotated with the closest gene, which is listed above the symbols in each locus.

**Table 2.** Overlapping loci between FTD and immune-mediated disease at a conjunction FDR < 0.05.

| SNP | Chr | Nearest<br>Gene | Associated<br>Phenotype | Associated<br>Phenotype<br>*p*-value | Min Conj<br>FDR | FTD<br>*p*-value | Direction<br>of Allelic<br>Effect |
| --- | --- | --- | --- | --- | --- | --- | --- |
| rs2269423 | 6 | *AGPAT1* | Ced | 4.63E-02 | 4.63E-02 | 6.28E-01 | +/- |
| rs3117097 | 6 | *BTNL2* | RA | 8.21E-05 | 8.21E-05 | 7.19E-03 | +/+ |
| rs204989 | 6 | *GPSM3* | UC | 2.90E-02 | 9.02E-03 | 2.58E-01 | +/+ |
| rs204991 | 6 | *GPSM3* | T1D | 1.00E-02 | 9.02E-03 | 2.58E-01 | +/+ |
| rs17427887 | 6 | *HLA-DQA2* | RA | 3.70E-02 | 3.70E-02 | 6.02E-01 | -/- |
| rs10484561 | 6 | *HLA-DQB1* | T1D | 1.86E-02 | 1.71E-02 | 4.17E-01 | -/+ |
| rs3135353 | 6 | *HLA-DRA* | CD | 2.29E-02 | 3.58E-03 | 1.35E-01 | +/+ |
| rs9268852 | 6 | *HLA-DRA* | UC | 1.25E-07 | 1.25E-07 | 1.03E-04 | +/+ |
| rs3129890 | 6 | *HLA-DRA* | T1D | 5.54E-05 | 5.54E-05 | 7.19E-03 | +/+ |
| rs6457590 | 6 | *HLA-DRA* | RA | 1.52E-02 | 1.52E-02 | 3.80E-01 | +/+ |
| rs9268877 | 6 | *HLA-DRA* | RA | 3.64E-07 | 1.25E-07 | 1.03E-04 | +/- |
| rs875142 | 6 | *PAQR8* | CD | 4.62E-02 | 4.62E-02 | 5.51E-01 | -/- |
| rs9261536 | 6 | *TRIM15* | T1D | 4.31E-02 | 4.31E-02 | 6.98E-01 | -/+ |
| rs3094138 | 6 | *TRIM26* | T1D | 4.63E-02 | 2.87E-02 | 6.28E-01 | -/+ |
| rs7778450 | 7 | *TNS3* | CeD | 4.26E-02 | 4.26E-02 | 6.17E-01 | -/- |
| rs2192493 | 7 | *TWISTNB* | T1D | 4.17E-02 | 4.17E-02 | 6.09E-01 | +/+ |
| rs10216900 | 8 | *CR590356* | T1D | 3.09E-02 | 3.09E-02 | 6.40E-01 | +/+ |
| rs10784359 | 12 | *SLC2A13* | RA | 3.33E-02 | 3.33E-02 | 5.90E-01 | +/+ |
| rs17572851 | 17 | *MAPT* | T1D | 2.47E-02 | 2.47E-02 | 5.90E-01 | +/+ |
| rs199533 | 17 | *NSF* | CD | 3.90E-02 | 1.95E-02 | 4.56E-01 | +/+ |
| rs2134297 | 18 | *DCC* | CeD | 3.11E-02 | 3.11E-02 | 5.73E-01 | +/+ |

Abbreviations: CeD, Celiac disease; Chr, Chromosome location; CD, Crohn disease;
FTD, Frontotemporal dementia; Min Conj FDR, minimum conjunction false discovery
rate; PSOR, Psoriasis; RA, Rheumatoid arthritis; SNP, Single-nucleotide polymorphism;
T1D, Type 1 diabetes; UC, Ulcerative colitis; –, negative effect estimate; +, positive effect estimates.

**Figure 2.**
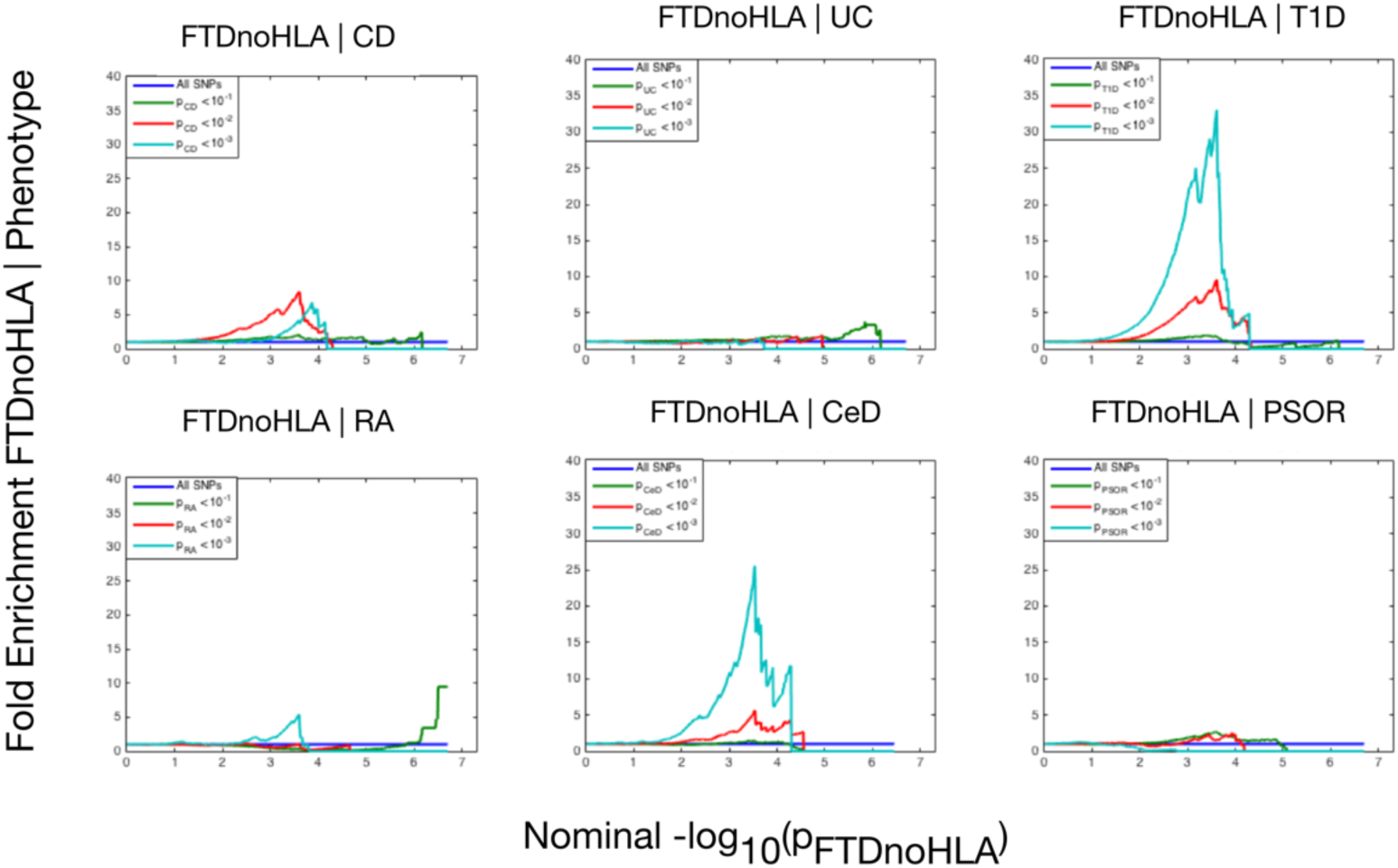
Fold enrichment plots of enrichment (after removing all regions in LD with *HLA* on chr 6) versus nominal -log_10_ *p*-values (corrected for inflation) in frontotemporal dementia (FTDnoMHC) below the standard GWAS threshold of *p* < 5× 10^-8^ as a function of significance of association with 6 immune-mediated diseases, namely Crohn’s disease (CD), ulcerative colitis (UC), type 1 diabetes (T1D), rheumatoid arthritis (RA), celiac disease (CeD), and psoriasis (PSOR) and at the level of -log10(*p*) ≥ 0, -log10(*p*) ≥ 1, -log10(*p*) ≥ 2 corresponding to *p* ≤ 1, *p* ≤ 0.1, *p* ≤ 0.01, respectively. Teal line indicates all SNPs.

Outside the *HLA* region, we found 7 other FTD-immune associated SNPs (Figure 2, Table 2), including two in strong LD, which mapped to the H1 haplotype of microtubule associated protein tau (*MAPT)* (LD: rs199533 with rs17572851; nearest genes: *NSF* and *MAPT*, pairwise D’ = 1, r^2^ = 0.94). Beyond *MAPT*, we found five additional novel loci associated with increased FTD risk, namely: 1) rs2192493 (chr 7, nearest gene = *TWISTNB*), 2) rs7778450 (chr 7, nearest gene = *TNS3*), 3) rs10216900 (chr 8, nearest gene = *CR590356*), 4) rs10784359 (chr 12, nearest gene = *SLC2A13*), and 5) rs2134297 (chr 18, nearest gene = *DCC*) (see Table 2 for additional details).

### Modest genetic enrichment between immune-mediated disease and PSP, CBD and ALS

To evaluate the specificity of the shared genetic overlap between FTD and immune-mediated disease, we also evaluated overlap between the 6 immune-mediated diseases and CBD, PSP, and ALS. For CBD and PSP a few of the immune-mediated diseases produced genetic enrichment comparable to that seen for FTD (Supplementary Figures 1A-C, Supplementary Tables 1-3). For example, we found 150-fold genetic enrichment between CBD and CeD, and 180-fold enrichment between PSP and RA. In contrast, we found minimal enrichment between ALS and the immune-mediated diseases tested, with the highest levels of enrichment between ALS and RA (up to 20-fold), and between ALS and CeD (up to 15-fold).

At a conjunction FDR< 0.05, we identified several SNPs associated with both CBD, PSP, or ALS and immune-mediated disease (Supplementary Figures 2A-C, Tables 1-3). Few of the SNPs shared between CBD, PSP, or ALS and immune-mediated disease mapped to the *HLA* region. Only two PSP-immune SNPs mapped to the region of *MLN* and *IRF4* on chromosome 6 and no CBD-and ALS-immune SNPs mapped to the *HLA* region (Supplementary Figures 2A-C, Tables 1-3).

Beyond the *HLA* region, we found several overlapping loci between the immune-mediated diseases and CBD, PSP and ALS (Supplementary Figures 2A-C, Tables 1-3). For PSP: 1) rs7642229 with CeD (chr 3, nearest gene = *XCR1*, FDR *p* = 1.74 × 10^-2^); 2) rs11718668 with CeD (chr 3, nearest gene = *TERC*, FDR *p* = 3.00× 10^-2^); 3) rs12203592 with CeD (chr 6, nearest gene = *IRF4*, FDR *p* = 4.17 × 10^-2^); 4) rs1122554 with RA (chr 6, nearest gene = *MLN*, FDR *p* = 2.09 × 10^-2^); 5) rs3748256 with RA (chr 11, nearest gene= *FAM76B*, FDR *p* = 2.09× 10^-2^). For ALS: 1) rs3828599 with CeD (chr 5, nearest gene= *GPX3*, FDR *p* = 2.27 × 10^-2^); 2) rs10488631 with RA (chr 7, nearest gene = *TNPO3*, FDR *p* = 3.42 × 10^-2^).

### cis-eQTL expression

To investigate whether shared FTD-immune SNPs modify gene expression, we evaluated *cis-*eQTLs in both brain and blood tissue types. At a previously established conservative Bonferroni corrected *p*-value < 3.9 ×10^-5^ [33], we found significant *cis*-associations between shared SNPs and genes in the *HLA* region on chromosome 6 in peripheral blood mononuclear cells (PBMC), lymphoblasts, and the human brain (see Supplementary Table 4 for gene expression associated with each SNP). We also found that rs199533 and rs17572851 on chr 17 were significantly associated with *MAPT* (*p* = 2 × 10^-12^) expression in the brain. Beyond the *HLA* and *MAPT* regions, notable cis-eQTLs included rs10784359 and *LRRK2* (*p* = 1.40 × 10^-7^) and rs2192493 and *TBKBP1* (*p* = 1.29 × 10^-6^) (see Supplementary Table 4).

### Protein-protein and co-expression networks

We found physical interaction and gene co-expression networks for the FTD-immune pleiotropic genes with significant *cis*-eQTLs (at a Bonferroni corrected p-value < 3.9 × 10^-5^). We found robust co-expression between various *HLA* classes further suggesting that large portions of the *HLA* region, rather than a few individual loci, may be involved with FTD (Fig 3, Supplementary Table 5). Interestingly, we found that several non-*HLA* functionally expressed FTD-immune genes, namely *LRRK2*, *PGBD5*, and *TBKPB1*, showed co-expression with *HLA* associated genes (Figure 3).

**Figure 3.**
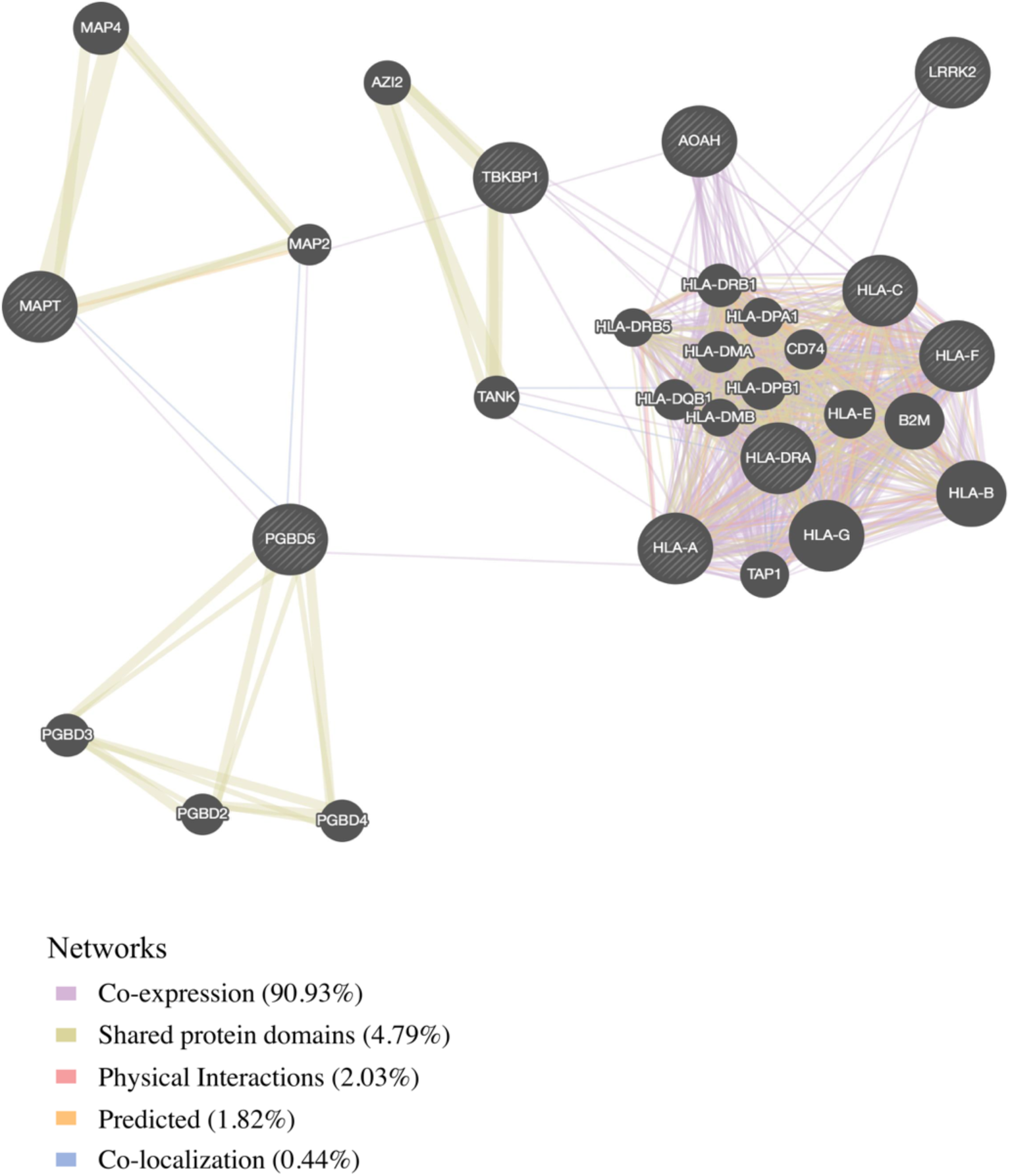
Network interaction graph predominantly illustrating co-expression and shared protein domains for functionally expressed FTD-immune pleiotropic genes.

### Genetic expression in FTD brains compared to controls

To investigate whether the FTD-immune pleiotropic genes with significant *cis*-eQTLs are differentially expressed in FTD brains, we compared gene expression in FTD-U brains to brains from neurologically healthy controls. We found significantly different levels of *HLA* gene expression in FTD-U brains compared to controls (Table 3). This was true of FTD-U brains regardless of progranulin gene (*GRN*) mutation status. In spite of the fact that the FTD GWAS used to identify these genes was based on patients with sporadic FTD (without *GRN* mutations), *GRN* mutation carriers showed the greatest differences in *HLA* gene expression (Figure 4, Table 3). These findings are compatible with prior work showing microglial-mediated immune dysfunction in *GRN* carriers [3,35].

**Table 3.** Genes associated with FTD and immune-mediated disease differentially altered in FTD-U patients versus controls.

| **Gene** | **Ctrl v. FTD-U *p*-value** | **Ctrl v. *GRN* *p*-value** |
| --- | --- | --- |
| *AOAH* | 0.631 | **0.013** |
| *HLA-A* | 0.622 | **0.004** |
| *HLA-C* | 0.372 | **0.002** |
| *HLA-F* | 0.592 | **0.001** |
| *HLA-DRA*<br>(*LOC101060835*) | **0.017** | **0.008** |
| *HLA-DRQ*<br>(*HLA-DQB1*) | **0.009** | **0.001** |
| *MAPT* | 0.376 | 0.090 |
| *TBKBP1* | **0.005** | 0.108 |
| *PGBD5* | 0.528 | **0.016** |
| *LRRK2* | N/A | N/A |

*P* values $\leq 0.05$ are in bold.

### Microglial enrichment

For the FTD-immune pleiotropic genes with significant *cis*-eQTLs, across different CNS cell types, we found significantly greater expression within microglia compared to neurons or fetal astrocytes (Figure 5A). Interestingly, *HLA* genes showed the greatest degree of differential expression. Four of the FTD-immune *HLA* associated genes, namely *HLA-DRA, AOAH, HLA-A,* and *HLA-C*, showed highest expression in microglia (ranging from 10 to 60 FPKM, see Figure 5B). In addition, *MAPT* was predominantly expressed in neurons, *LRRK2* in microglia/macrophages, *PGBD5* in neurons, and *TBKBP1* in fetal astrocytes (Figure 5B and Supplemental Figures 3A-C).

**Figure 4.**
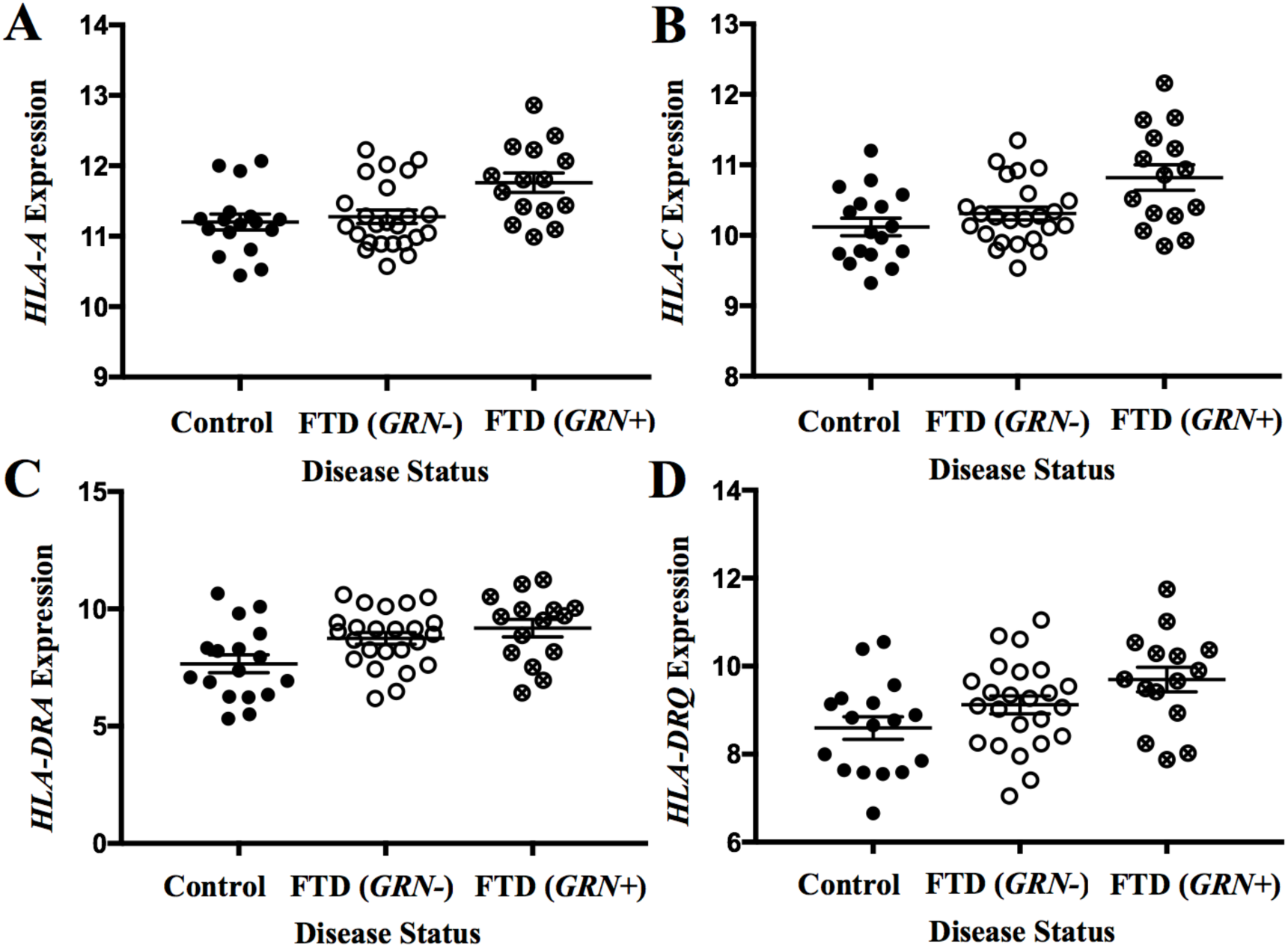
FTD-Immune genes are elevated in FTD-*GRN* brains. Expression for the genes with the largest effect sizes are plotted: A) *HLA-A*, B) *HLA-C*, C) *HLA-DRA*, and D) *HLA-DRQ*. Expression values were obtained from GSE13162 for FTD-U brains with and without *GRN* mutations and neuropathology-free controls. Horizontal bar represents mean ± SEM.

## DISCUSSION

We systematically assessed genetic overlap between 4 FTD-related disorders and several immune-mediated diseases. We found extensive genetic overlap between FTD and immune-mediated disease particularly within the *HLA* region on chromosome 6, a region rich in genes associated with microglial function. This genetic enrichment was specific to FTD and did not extend to CBD, PSP, or ALS. Further, we found that shared FTD-immune gene variants were differentially expressed in FTD patients compared with controls, and in microglia compared with other CNS cells. Beyond the *HLA* region, by leveraging the immune-mediated traits, we detected novel FTD susceptibility loci within *LRRK2, TBKBP1* and *PGBD5.* Considered together, these findings suggest that various microglia and inflammation-associated genes, particularly within the *HLA* region, play a critical and selective role in FTD pathogenesis.

By combining GWAS from multiple studies and applying a pleiotropic approach, we identified genetic variants jointly associated with FTD-related disorders and immune-mediated disease. We found that the strength of genetic overlap with immune-mediated disease varies markedly across FTD-related disorders, with the strongest pleiotropic effects associated with FTD, followed by CBD and PSP, and the weakest pleiotropic effects associated with ALS. We identified eleven FTD-immune associated loci that mapped to the *HLA* region, a concentration of loci that was not observed for the other disorders. Indeed, only two PSP-immune pleiotropic SNPs and no CBD-or ALS-immune pleiotropic SNPs mapped to the *HLA* region. Previous work has identified particular *HLA* genes associated with CBD, PSP, and ALS [34,36]. In contrast, our current findings implicate large portions of the *HLA* region in the pathogenesis of FTD. Together these results suggest that each disorder across the larger FTD spectrum has a unique relationship with the *HLA* region.

Our results also indicate that functionally expressed FTD-immune shared genetic variants are differentially expressed in FTD brains compared to controls and in microglia compared to other CNS cell types (Figure 5). Microglia play a role in the pathophysiology of *GRN*+ FTD. Progranulin is expressed in microglia [35] and *GRN* haploinsufficiency is associated with abnormal microglia activation and neurodegeneration [3]. It is perhaps expected, therefore, that *GRN+* brains show differential expression of FTD-immune genes, even though these genetic variants were derived from GWAS of patients with sporadic FTD (who lack *GRN* or other established FTD mutations). More surprising is the presence of similar enrichment in *GRN*-brains,suggesting that dysfunction of microglial-centered immune networks may be a common feature of FTD pathogenesis.

**Figure 5.**
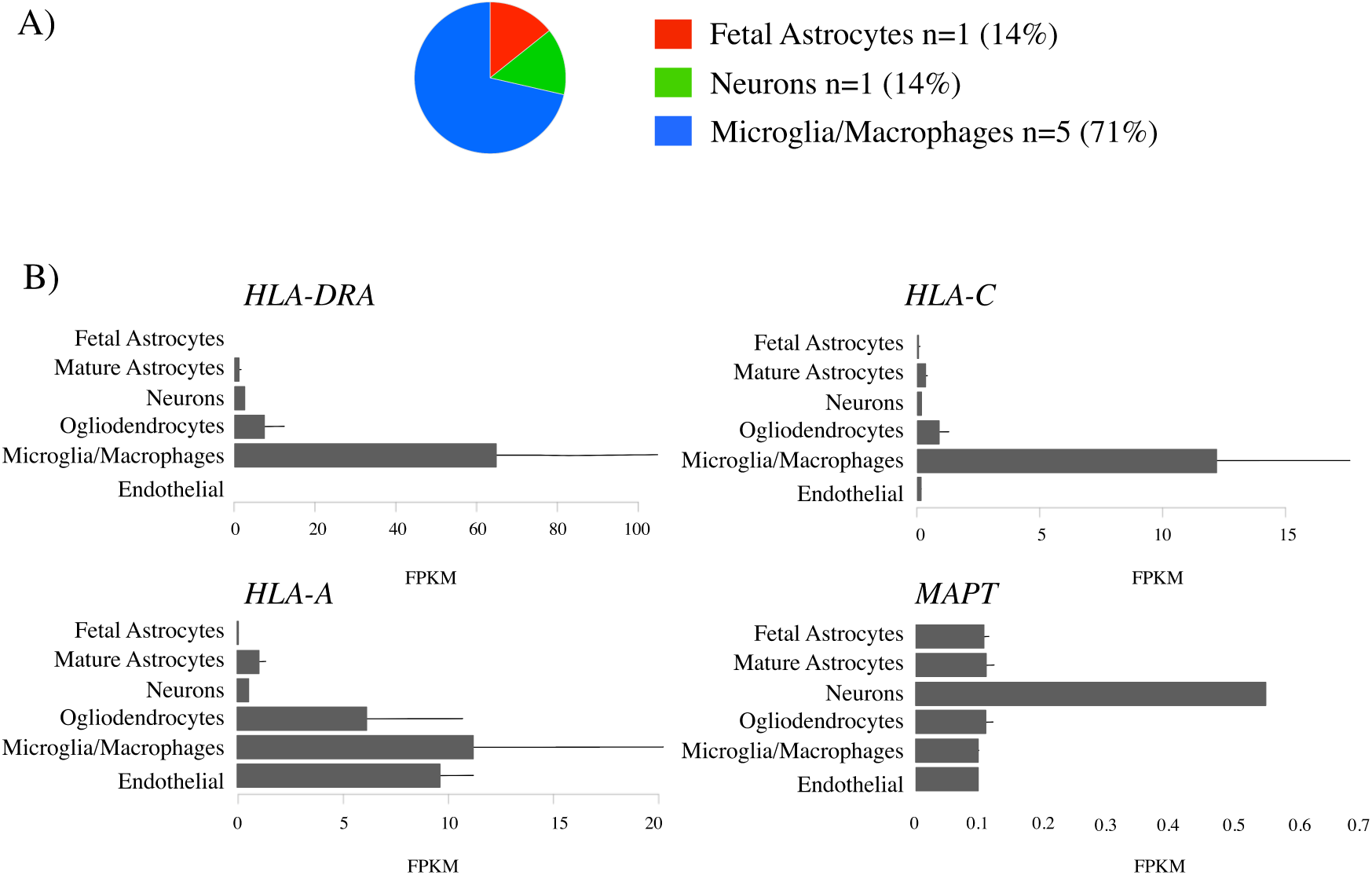
Microglia enrichment in FTD-Immune genes. FTD-Immune genes were analyzed to determine the cell-type in which each gene is most highly expressed [32]. A) Pie chart plotting the relative number of genes most highly expressed in in each cell type. No genes were most highly expressed in endothelial cells or oligodendrocytes. B) Individual bar plots showing cell-type specific expression for genes with the largest effect size.

By leveraging statistical power from the large immune-mediated GWASs, we identified novel FTD susceptibility loci within *LRRK2, TBKBP1* and *PGBD5* and confirmed previously shown FTD associated signal within the *MAPT* region*. LRRK2* mutations are a cause of Parkinson's disease [37] and Crohn’s disease [38]. We previously described a potential link between FTD and the *LRRK2* locus [39] and another study using a small sample showed that *LRRK2* mutations may increase FTD risk [40].Together these results suggest that the extended *LRRK2* locus might influence FTD despite common genetic variants within *LRRK2* not reaching genome-wide significance in large FTD-GWAS [5]. We suggest that increased expression of *LRRK2* in microglia results in proinflammatory responses, possibly by modulating TNF-alpha secretion (Tumor-Necrosis Factor) [41]. *TBKBP1* also modulates TNF-alpha signaling by binding to *TBK1* (*TANK*-binding kinase 1) [42]; rare pathogenic variants in *TBK1* cause FTD-ALS [43,44,45]. Importantly, elevated CSF levels of TNF-alpha are a core feature of FTD [6,46]. Building on these findings, in our bioinformatics ‘network’ based analysis, we found that both *LRRK2* and *TBKBP1* interact with genes within the *HLA* region (Figure 3). Further, physical interactions between *MAPT* and the *HLA* network are compatible with research suggesting that under different conditions reactive microglia can either drive or mitigate tau pathology [47,48]. *MAPT* mutations, which disrupt the normal binding of tau protein to tubulin, account for a large proportion of familial FTD cases [49]. Together, these findings suggest that *LRRK2, TBKBP1,* and *MAPT* may, at least in part, influence FTD pathogenesis via *HLA*-related mechanisms.

This study has limitations. Particularly, in the original datasets that form the basis of our analysis, diagnosis of FTD was established clinically. Given the clinical overlap among neurodegenerative diseases, we cannot exclude the potential influence of clinical misdiagnosis. As such, our findings would benefit from confirmation in large pathologically confirmed cohorts. In addition, given the complex interconnectedness of the *HLA* region (see Figure 3), we were not able to define the specific gene(s) on chromosome 6 responsible for our pleiotropic signal. However, given the number of *HLA* loci associated with increased FTD risk, it may be the case that no single *HLA* variant will be clinically informative; rather, an *additive combination* of these microglia-associated genetic variants may better inform FTD risk.

In conclusion, across a large cohort (total n = 192,886 cases and controls), we leveraged pleiotropy between FTD-related disorders and immune-mediated disease to identify several genes within the *HLA* region that are expressed within microglia and differentially expressed in the brains of patients with FTD. Building on prior work [6,7], our results suggest that immune dysfunction is central to the pathophysiology of a subset of FTD patients. These findings have important implications for future work focused on monitoring microglial activation as a marker of disease progression and on developing anti-inflammatory therapies to modify disease outcomes in patients with FTD.

## ACKNOWLEDGEMENTS

Primary support for data analyses was provided by AG046374 (CMK), U24DA041123 (AMD, LPS, RSD), National Alzheimer's Coordinating Center (NACC) Junior Investigator (JI) Award (RSD), RSNA Resident/Fellow Grant (RSD), Foundation of ASNR Alzheimer’s Imaging Grant (RSD), Alzheimer’s Society Grant 284 (RF), ARRS/ASNR Scholar Award (LPS), and the Tau Consortium. Additional support was provided by the Larry L. Hillblom Foundation 2016-A-005-SUP (JSY), AFTD Susan Marcus Memorial Fund Clinical Research Grant (JSY), NIA K01 AG049152 (JSY), P01-AG-017586 (GDS), the Bluefield Project to Cure FTD (JSY), and the Tau Consortium (JSY, GDR). GUH was supported by the Deutsche Forschungsgemeinschaft (DFG, HO2402/6-2 & Munich Cluster for Systems Neurology SyNergy), the German Federal Ministry of Education and Research (BMBF, 01KU1403A), and the NOMIS Foundation (FTD project). The PSP-GWAS was funded by a grant from the CurePSP Foundation, the Peebler PSP Research Foundation.

